# Evolution of predator foraging in response to prey infection favors species coexistence

**DOI:** 10.1101/2020.04.18.047811

**Authors:** Loïc Prosnier, Vincent Médoc, Nicolas Loeuille

**Affiliations:** Sorbonne Université, Université de Paris, Université Paris-Est Créteil, CNRS, INRAE, IRD, institute of Ecology and Environmental Science of Paris (iEES-Paris), 75005 Paris, France; CItés, TERritoires, Environnement et Sociétés (CITERES, CNRS UMR 7324), University of Tours, 37200 Tours, France; Equipe de Neuro-Ethologie Sensorielle (ENES), Centre de Recherche en Neurosciences de Lyon (CRNL), CNRS, INSERM, Universite de Lyon/Saint-Etienne, 42100 Saint-Etienne, France

**Keywords:** Adaptive foraging, virulence, vulnerability, predator diet, profitability, parasitism

## Abstract

As acknowledged by optimal foraging theory, predator diets depend on prey profitability. Parasites, affecting prey within food webs, are known to affect simultaneously host vulnerability to predation and host energy contents, thereby affecting profitability. In this work, we study the eco-evolutionary consequences of prey infection by a non trophically-transmitted parasite, with a simple lifecycle, on predator diet. We also analyse the consequences for coexistence between prey, predators and parasites. We model a trophic module with one predator and two prey species, one of these prey being infected by a parasite, and distinguish two effects of infection: a decrease in host fecundity (virulence effect) and an increase in vulnerability to predation (facilitation effect). Predator foraging may evolve toward specialist or generalist strategies, the latter being less efficient on a given resource. We show that virulence effects lead to specialisation on the non-infected prey while facilitation effects, by increasing prey profitability, favor specialisation on the infected prey. Combining the two effects at intermediate intensities promotes either generalist predators or the diversification of foraging strategies (coexistence of specialists), depending on the trade-off shape. We then investigate how the evolution of predator diet affects the niche overlap between predator and parasite. Facilitation effects systematically lead to a high niche overlap, ultimately resulting in the loss of the parasite. Virulence effects conversely favor coexistence by allowing a separation of the predator and parasite niches.

## 1. Introduction

Parasitism is an important but often overlooked part of food webs. When assessed, parasites make for a large part of local biomass (Kuris et al., 2008) and substantially increase the number of interactions within the ecological network (Amundsen et al., 2009). Consequently, there is increasing research about the consequences of parasitism in terms of diversity (Hatcher et al., 2006, 2008; Hudson et al., 2006), stability (Hilker & Schmitz, 2008; Hudson et al., 1998), and energy fluxes (Buck & Ripple, 2017; Sato et al., 2012). Many works highlight that predators affect parasites (Richards et al., 2022), for instance when parasites serve as concomitant prey (Cirtwill & Stouffer, 2015; Hatcher et al., 2008; Johnson et al., 2010), or when predation promotes the spread of parasites (Cáceres et al., 2009; Richards et al., 2023). However, in such complex systems, it remains unclear how the interplay of parasitism and predation affects coexistence (Lopez & Duffy, 2021). Effects on evolutionary dynamics are equally unclear. Most works indeed focus on host-parasite evolution in the absence of trophic contexts (Auld et al., 2013; Choo et al., 2003), despite the ecological importance of the evolution of predator diet for coexistence and stability (Kondoh, 2003). Consequently, we ask in this work how parasites could affect the evolution of predator diet, in particular its niche specialisation (Britton & Andreou, 2016), and ultimately the ecological repercussions of such evolutionary dynamics.

Infection has two main effects on host phenotype and reproductive success that must be considered. First, a direct consequence of infection can be a reduction of host fecundity (Decaestecker et al., 2003; Hurd, 2001) and/or survival (Brassard et al., 1982; Lynsdale et al., 2017; Prins & Weyerhaeuser, 1987), what we called “virulence effect” hereafter. Second, indirect effects mediated by the trophic context can occur when alterations in the behaviour, morphology or physiology of hosts increase their vulnerability to predation (Goren & Ben-Ami, 2017; Hudson et al., 1998; Wale et al., 2021). We use the term “facilitation effect” hereafter. Virulence and facilitation effects can be a byproduct of energy reallocation within the host to the benefit of the parasite (Forshay et al., 2008) often occur simultaneously. For instance, researchers have analysed the infection of *Daphnia magna* by Daphnia Iridovirus 1 (DIV-1). DIV-1 was known to negatively affect both fecundity and mortality (Ebert et al., 2000). More recently, Prosnier et al. (2024) observed a decrease of host survival and mobility, and an increase of host coloration, size, and energy content (due to higher protein content). Handling time by a predatory arthropod (*Notonecta sp.*) was also increased. DIV-1 infection therefore incurs a strong virulence effect (reduction of survival by 2/3) and a facilitation effect that increases host profitability by 24%.

From an ecological point of view, Prosnier et al. (2018) explored the implications of the virulence and facilitation effects (interaction effect in their paper) within an eco-epidemiological model that includes a predator and two prey species with one of them infected by parasites, evolution being ignored. Their results suggest that ecological coexistence is possible for intermediate levels of virulence and facilitation effects when parasites affect the most competitive prey. Prey coexistence is here explained by the balance between direct and apparent competition (Gause, 1934; Holt, 1977; Holt et al., 1994), respectively influenced by the virulence and facilitation effects. Competition also exists between the parasite and the predator. Higher virulence reduces predator resources, while facilitation favors predator and reduces parasite prevalence as in the healthy herd hypothesis (Duffy et al., 2005; Lafferty, 2004; Packer et al., 2003), this effect increasing with the consumption of infected individuals (Brandell et al., 2022).

While Prosnier et al (2018) therefore suggests that coexistence of predators and parasites requires specific conditions, it is widely reported in food webs (Byers, 2009; Kagami et al., 2014; Lafferty et al., 2008). Accounting for evolutionary changes in predator diets based on the principle of adaptive foraging – known to promote coexistence and stability in complex networks (Kondoh, 2003) – could explain coexistence in the presence of parasites. Under optimal foraging theory, predators are assumed to interact only with prey species that enhance their net energy intake rate (Pyke et al., 1977). The relative profitability of prey species, defined as the ratio between energetic gain (prey energetic value) and energetic cost (search and handling times), determines the occurrence of trophic links and therefore the degree of generalism among predators (Charnov, 1976a, 1976b; Emlen, 1966; MacArthur & Pianka, 1966). Whenever parasitism enhances prey profitability (as in the DIV1-*Daphnia magna* system), evolution should therefore lead to specialisation on infected individuals. However, parasites can also drastically decrease prey abundances due to high virulence, leading to a low availability of infected prey and favoring the attack of alternative (i.e., non-infected) prey. Consequently, coexistence could be increased if diet evolution under virulence effects leads to niche separation between the predator and the parasite, or decreased if facilitation effects promote niche convergence.

To investigate these eco-evolutionary dynamics, we modelled a trophic module with one predator and two prey in competition, the most competitive prey being infected by a parasite and structured into susceptible and infected individuals. The parasite, which has a simple life cycle and a horizontal transmission, induces virulence and facilitation effects modelled respectively by a reduction of host fecundity and an increase in host vulnerability to predation as in Prosnier et al. (2018). We study the evolution of the predator’s trophic niche, assuming a trade-off between the consumption of the two prey species. We consider flexible trade-off shapes, higher trade-off concavity usually favoring generalist strategies (Egas et al., 2004; Rueffler et al., 2006). In agreement with our predictions, we show that virulence effects favor specialisation on the alternative prey while facilitation effects have the opposite effect. Our results also show that evolution favors niche separation among parasites and predators in case of a virulent parasite, thereby promoting coexistence.

## 2. Model presentation and analysis

### 2.1. Predation-infection model

The model considers two prey species sharing a predator. We assume both intraspecific and interspecific competition among prey and we model predation through linear functional responses (Novak et al., 2025). The infected prey (labelled 1) is also the most competitive and its population is structured into susceptible and infected individuals (*S_1_* and *I_1_* respectively). Parasite transmission is horizontal, and we assume that infected individuals do not recover. Considering these hypotheses, ecological dynamics follow the set of equations:

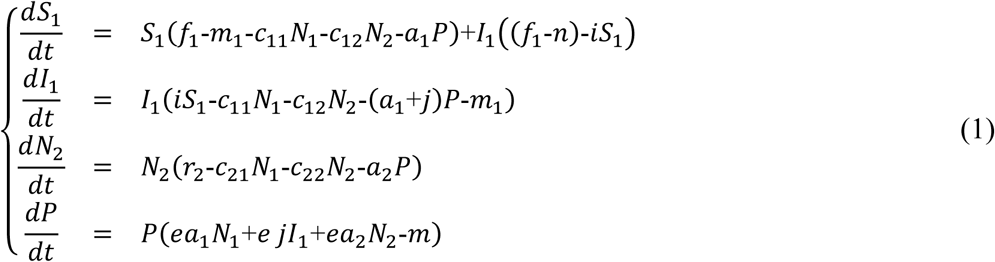

with *N_x_* the total prey species *x* population, *S_x_* and *I_x_* respectively the susceptible and infected individuals (thus *N*_1_=*S*_1_+*I*_1_), *f*_1_ the intrinsic fecundity rate of the infected prey, *m*_1_ its intrinsic mortality rate, *r*_2_ is the intrinsic growth rate of the alternative prey (with *r*_2_=*f*_2_-*m*_2_), *P* the predator density, *m* its intrinsic mortality rate, *i* the *per capita* transmission rate of the parasite, *c_xx_* the *per capita* competition rate of prey *x*, *c_xy_* the *per capita* competition effect of prey *y* on prey *x*, *a_x_* the *per capita* attack rate of the predator on prey *x*, *e* the conversion efficiency. While we acknowledge that infection could influence all prey parameters, we here focus on two changes. Virulence effects are implemented through a decreased fecundity (parameter *n*) and facilitation effects through an increased attack rate by the predator (parameter *j*). The biological interpretation, dimensions and default values of parameters are given in Table 1.

**Table 1.**
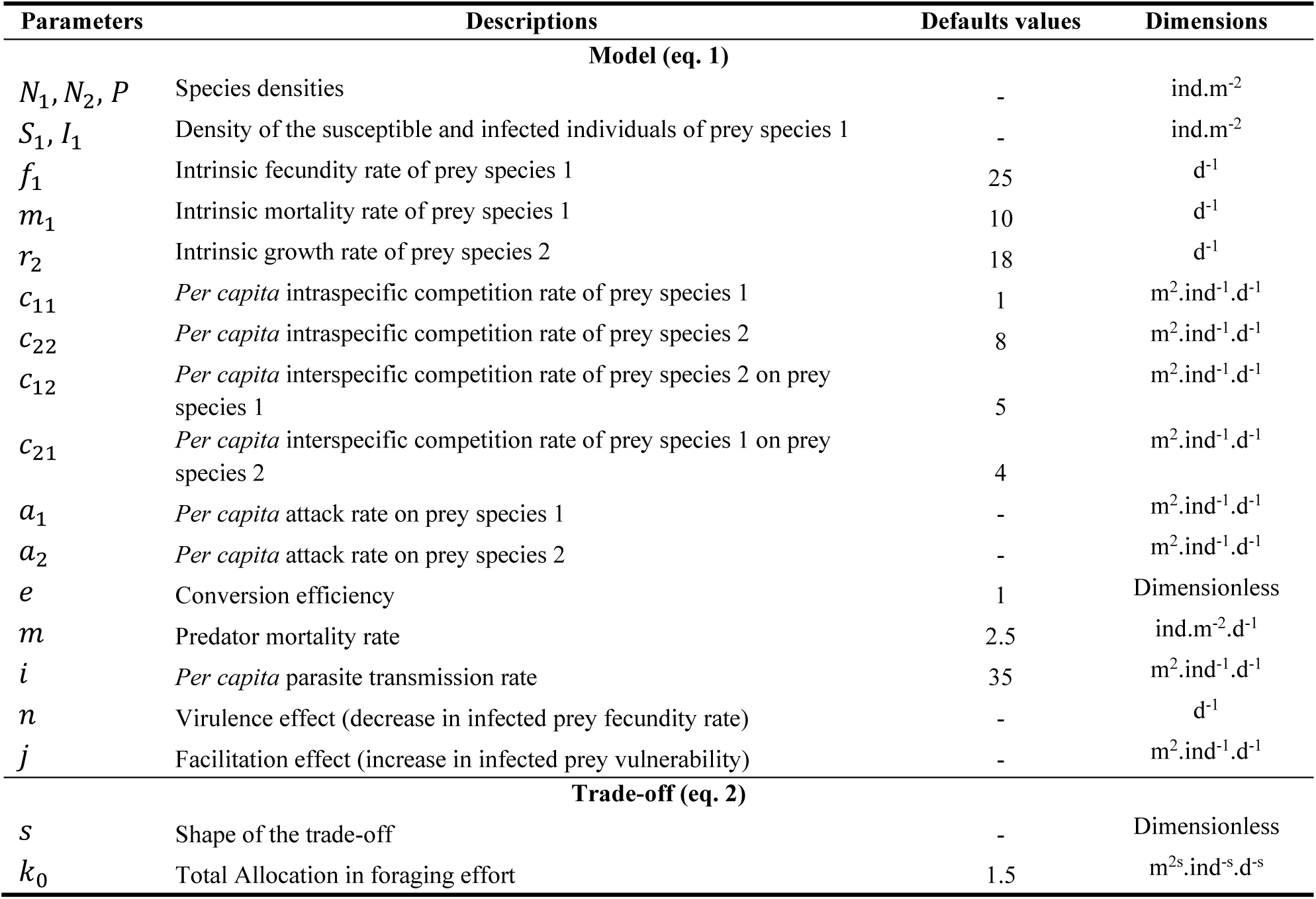
Biological interpretation, dimensions and default values of model’s parameters. Used values are those proposed in Prosnier et al. (2018) and based on Hutson and Vickers (1983).

Prey *N_1_* being the most competitive, we assume that *r*_1_*c*_22_>*r*_2_*c*_12_ and *r*_1_*c*_21_>*r*_2_*c*_11_ (Hutson & Vickers, 1983) so that it excludes the alternative prey in the absence of predators and parasites.

Because *r*_1_=*f*_1_-*m*_1_, the virulence effect (reduced fecundity), directly affects the competitive hierarchy, so that in the absence of predators, parasites reduce the advantage of the dominant competitor and favor coexistence. This situation, where coexistence is highly constrained without enemies, allows to better highlight the role of predators and parasites in the global coexistence.

A former analysis of the ecological model (Prosnier et al., 2018) showed that coexistence is possible only under restrictive conditions, and requires both virulence and facilitation effects, which, respectively, reduce the competitive ability of the dominant prey and make it the predator’s favorite prey. Too strong virulence effects allow the parasite to competitively exclude the predator, while too strong facilitation effects allow the predator to overconsume infected prey, thereby leading to parasite’s extinction.

### 2.2. Trade-off constraints on adaptive foraging

We assume that the foraging strategy of the predator varies along a trade-off function that describes allocation (e.g., attack rate) between the two prey species. The time or energy devoted to the consumption of one prey reduces predation on the other. We also assume that generalist strategies have lower attack rates on a given prey compared to a specialist of the same prey. Such trade-offs (Fig. 1) can be modelled using the following function (Egas et al., 2004):

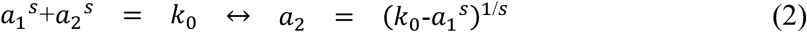

where *s* affects the trade-off curvature (convex: *s*<1, linear: *s*=1 or concave: *s*>1), and *k*_0_ corresponds to the total allocation of energy or time in predation.

**Figure 1.**
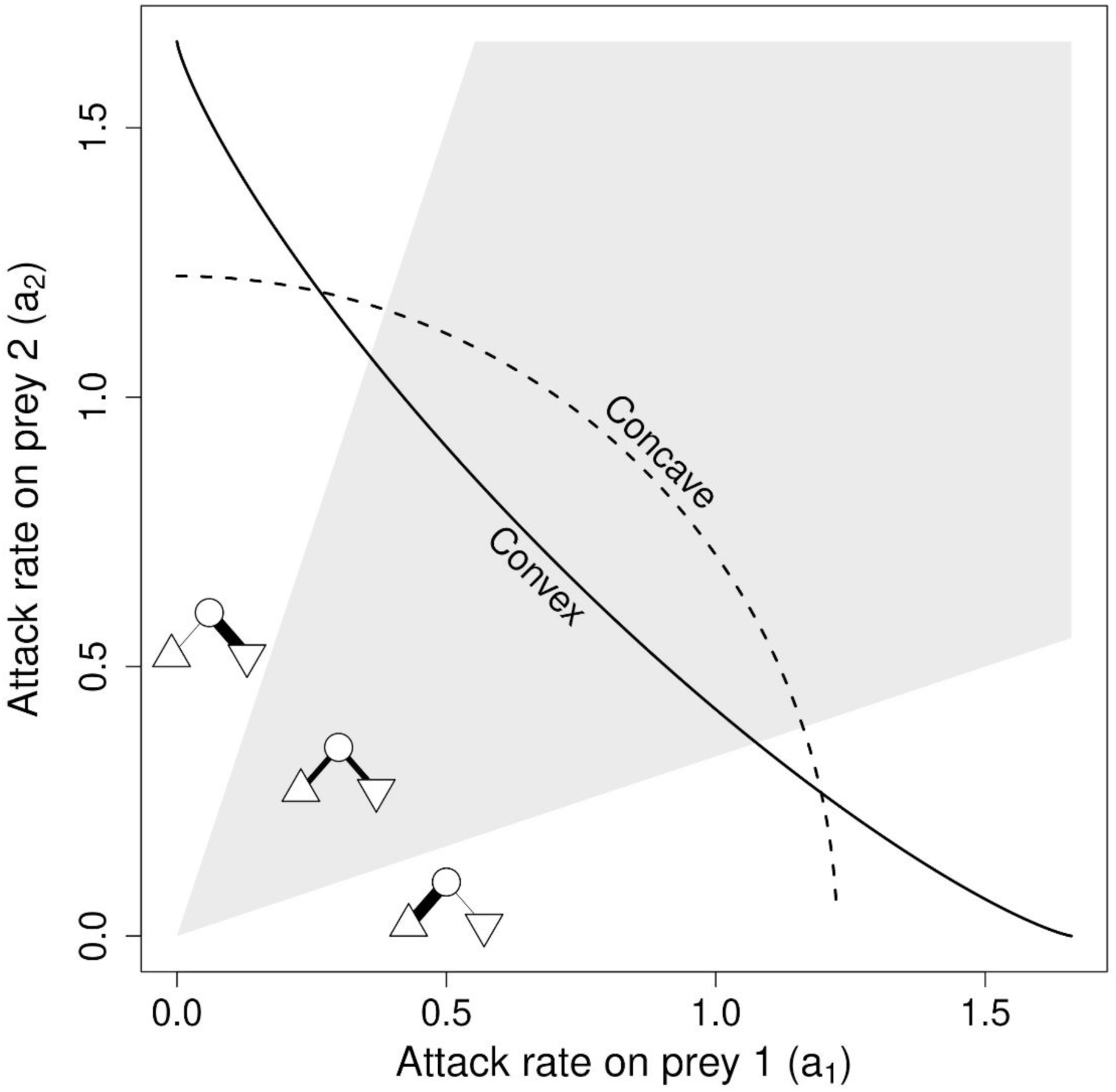
Concave (s>1) and convex (s<1) trade-off of predation investment in each prey. Modules display the intensity of each trophic link. Symbols represent modules composition: predator (circle), competitive/infected prey (triangle), alternative prey (inversed triangle). Bold lines show values for generalist predators and thin lines specialist predators. See Table 1 for parameter values.

### 2.3. Evolution of predator’s diet

We use adaptive dynamics (Dieckmann & Law, 1996; Geritz et al., 1998) to analyse diet evolution (i.e., variations in *a*_1_). Adaptive dynamics assume that ecological dynamics occur on faster timescales compared to evolution (i.e., rare mutations, of small phenotypic effects). Under such conditions, resident phenotypes *a*_1_ reach the ecological equilibrium before the next mutation occurs and evolution can be understood based on the invasibility of this equilibrium by nearby mutants (phenotype *a*_1*m*_). The relative fitness of mutants *ω*(*a*_1*m*_, *a*_1_) is defined by their intrinsic growth rate when rare in these equilibrium conditions (invasion fitness, Metz et al. (1992)). Given equation (1), the intrinsic growth rate of the mutant 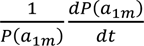 when mutants are rare can be written as:

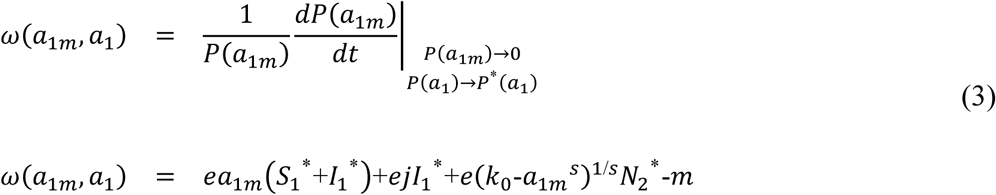

From equation (3), we see that the mutants having high attack rates on prey *N*_1_ (*a*_1_*_m_*) can invade (i.e., the fitness function is more likely positive) when *N*_1_ population is large enough, or when the density of the alternative prey *N*_2_ is small enough.

Variations of trait *a*_1_ (with *a*_2_ deduced using eq. 2), can be described using the canonical equation of adaptive dynamics (Dieckmann & Law, 1996):

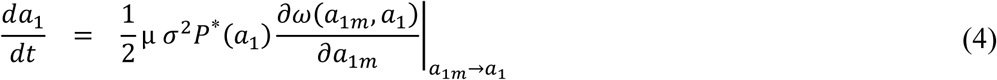

Where *µ* is the *per capita* mutation rate, *σ^2^*the phenotypic variance linked to mutation process, and *P\** the equilibrium density of residents (of phenotype *a*_1_). These three parameters affect the speed of evolution – trait evolves faster if mutation rates and amplitudes are high or if the resident population is large. In eq. 4, the selection process is described by the slope of the fitness landscape, 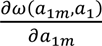, around the resident phenotype, a_1m_ → a_1_. The direction of evolution entirely depends on the sign of this slope named gradient fitness, 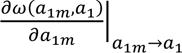 So that *a*_1_ increases when the slope is positive and decreases when the slope is negative.

From equation (4), note that phenotypes no longer vary when the gradient is null. This particular point is named evolutionary singularity, written 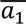. It satisfies:

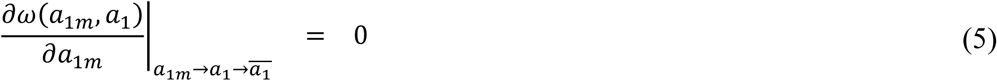

Dynamics close to the singularities can be characterized using two criteria (Diekmann, 2004; Eshel, 1983; Geritz et al., 1997): invasibility and convergence. Invasibility describes whether the singularity can be invaded by nearby mutants (i.e., non-invasible strategies are named Evolutionary Stable Strategy, ESS). Convergence describes if, starting close to the singularity, selected mutants are even closer to it, so that the strategy is eventually reached. Understanding convergence and invasibility is important, as it uncovers the overall dynamics, as analysed below. For instance, if the singularity is convergent and non-invasible, then it is called a continuously stable strategy (Eshel, 1983) and evolution eventually stops at this strategy. Conversely a convergent and invasible singularity would lead to the emergence of polymorphism (branching events, see below).

From Diekmann (2004), seconds derivatives of the fitness *ω*(*a*_1*m*_, *a*_1_) allow us to characterize invasibility and convergency. A singularity is non-invasible if:

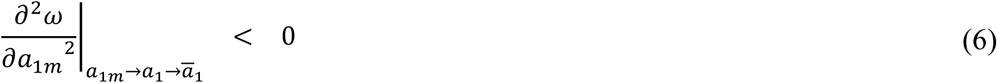

Thus if

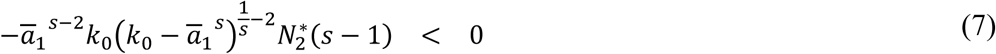

From this equation we can see that non-invasible singularities only occur with a concave trade-off (s>1).

A singularity is convergent if:

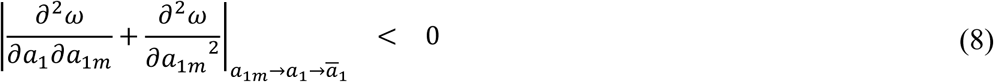

Thus if:

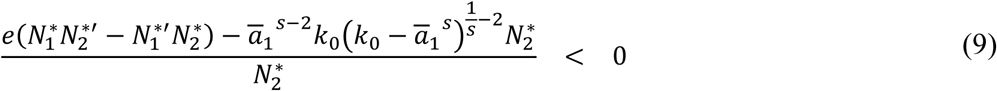

Each singularity *a*_1_^_^ corresponds to a certain level of foraging generalism. Assuming convergence, low values indicate that the predator mostly feeds on the alternative resource, while high values mean that the predator mainly feeds on the host. We consider a predator as a specialist if more than 75% of its diet is based on one prey (white areas on Fig. 1).

### 2.4. Various outcomes of diet evolution

In our system, the predator specialises on the most competitive prey (*N_1_*) in the absence of parasite (not shown). In the presence of parasites, we can observe four outcomes depicted on Figures 2, 3 and A1. Figure 2 represents the temporal dynamics of the four populations (uninfected prey, infected prey, non-host prey and predator) and the values of the attack rate on *a*_1_ for four different scenarios of virulence vs facilitation effects. The simulations were performed with Scilab (version 5.5.2). Figures 3 and A1 are Pairwise Invasibility Plots (PIP, Diekmann (2004)) – generated using Mathematica® 11.1.1 (Wolfram research). Such PIP plots show the sign of the relative fitness *ω*(*a*_1_*_m_*, *a*_1_) for different mutations (along the y-axis) given various resident populations (along the x-axis). The mutant has a positive relative fitness and thus can invade the population in the dark area, and a negative relative fitness in the white area. When mutations are small and rare, resident and mutant trait values are typically close to the diagonal *x*=*y*, and thus trait evolution follows the diagonal, depending on fitness value above and below the diagonal. We here plot these PIPs for increasing virulence effects (from left to right) and for concave and convex trade-offs. Evolutionary singularities are at the intersection of the diagonal and the other isocline. Their convergence and invasibility can be analysed from the relative position of black (positive fitness) and white (negative fitness) areas, which allows to deduce the general evolutionary dynamics (the direction of evolution is shown using white arrows).

**Figure 2.**
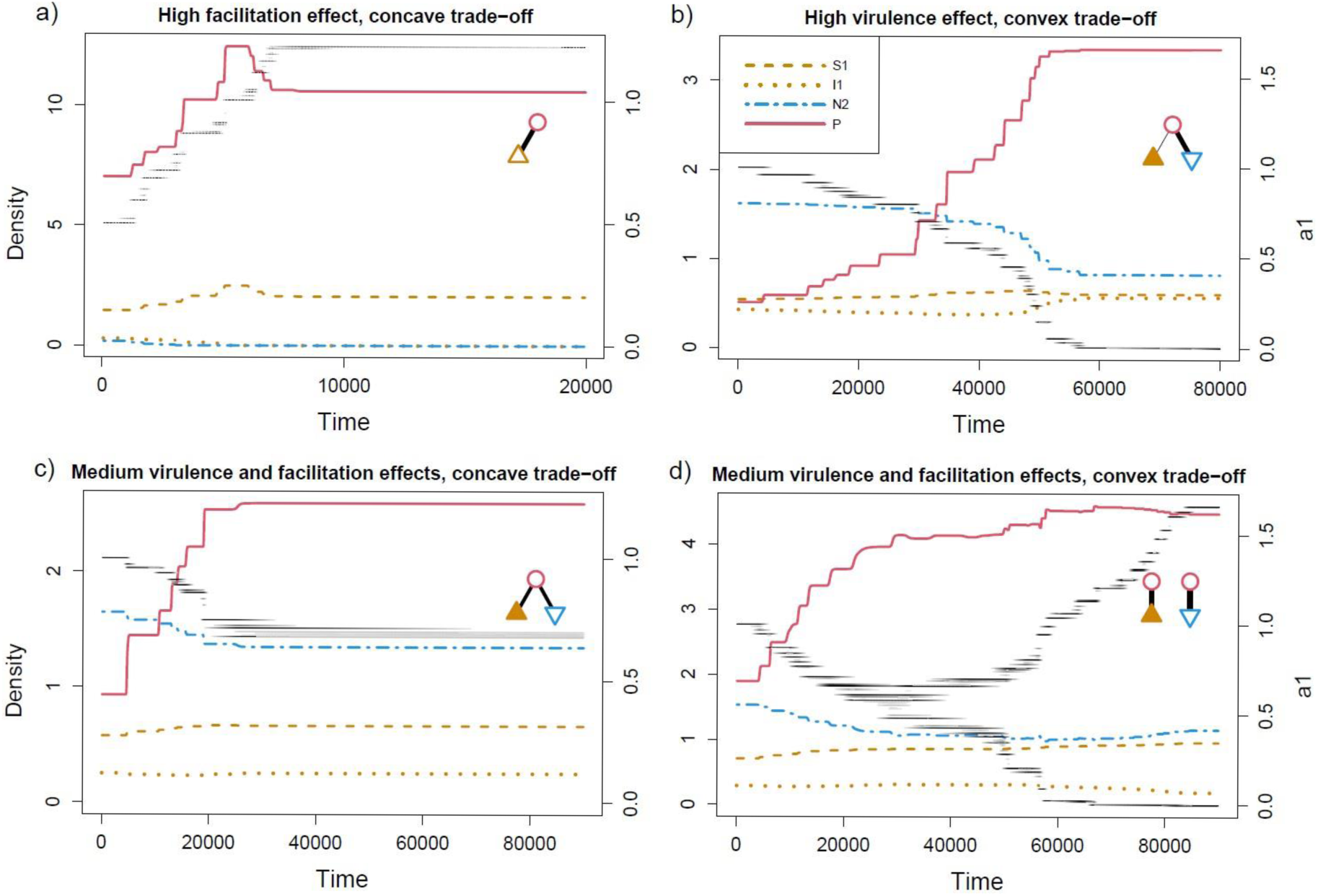
Various eco-evolutionary dynamics (species densities and trait value of the predator) of the module depending of trade-off shape and parasitism effects. a,c) concave trade-off, b,d) convex trade-off. a) high facilitation effect, b) high virulence effect, c,d) medium virulence and facilitation effects. Symbols show module’s composition and the intensity of trophic links at the end of the simulation: predator (circle), competitive prey (triangle), alternative prey (inversed triangle). Infected prey are shown in fdl and alternative prey in empty. Within infected prey, we display the density of susceptible individuals *S_1_* in orange dashed line and that of infected individuals *I_1_* in orange dotted line. The alternative prey *N_2_* is in blue dashed-dotted line and the predator *P* in red solid line. Values of *a_1_,* the evolving trait, are shown in dark (grey scale indicates the density of the predator for each trait value). Parameter values: see Table 1, except for *n,* the virulence effect, and *j,* the facilitation effect: a) *n* = 20,*j* = 5, *s* = 2; b) *n* = 15,*j* = 2, *s* = 0.8; c) *n* = 20,*j* = 2, *s* = 2; d) *n* = 15,*j* = 3, s = 0.8.

**Figure 3.**
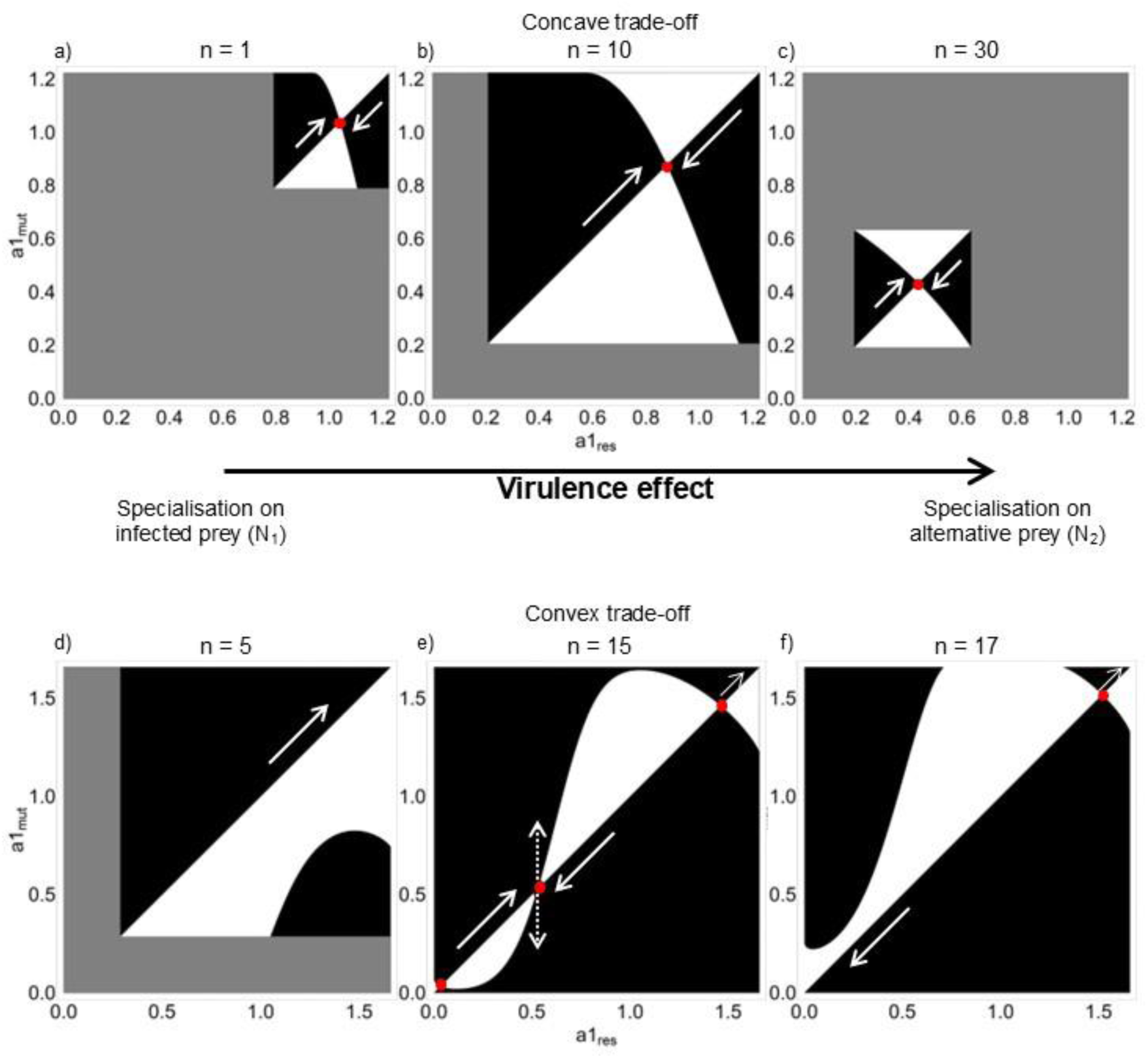
Pairwise Invasibility Plots showing the evolution of predator diet (i.e., evolution of *a_1_*) for concave (a-c) and convex (d-f) trade-offs, when increasing the intensity of the virulence effect, *n*. Black areas correspond to a positive relative fitness of mutants, white areas to a negative relative fitness of mutants, and grey areas shows no-coexistence of the three species. The white solid arrows show the direction of evolution, the white dotted arrows show evolutionary branching. a-c) Continuously Stable Strategy, d) no singularity, e) one Evolutionary Branching Points and two repellors, f) one repellor. Parameter values: see Table 1, except a) *n* = 1, *j* = 1, *s* = 2; b) *n* = 10, *j* = 1, *s* = 2; c) *n* = 30, *j* = 1, *s* = 2; d) *n* = 5, *j* = 3, *s* = 0.8; e) *n* = 15, *j* = 3, *s* = 0.8; f) *n* = 17, *j* = 3, *s* = 0.8. Note that Fig A1 shows the results with the facilitation effect.

Generally, we observe a specialisation on the infected prey when facilitation effects are high and a specialisation on alternative prey under high virulence effects. At intermediate effects, we obtain generalist predators with a concave trade-off, and the emergence of polymorphism under a convex trade-off. In detail, the predator can be generalist (Figs 2c, 3b, A1b,c) when the singularity *a*_1_^_^ is a Continuously Stable Strategy (CSS, i.e., convergent and non-invasible) of intermediate value (see figure 1). As previously showed (eq. 7), this only occurs for concave trade-offs. Predators can also evolve to become specialists, either on the host (Fig. 2a) or on the alternative prey (Fig. 2b), which can be achieved in four ways. First, specialists could correspond to CSS of large or small values (Fig. 3a,c, A1a). Alternatively, the singularity may be a Repellor (i.e., non-convergent and invasible). In such conditions, evolutionary trajectories move away from the singularity, leading to specialisation on one species or another depending on the initial diet. In our model, repellors only occur with convex trade-offs as they are invasible (eq. 7) (Fig. 3f). A third possibility to obtain a specialist is when no singularity exists, so that the fitness gradient (eq. 3) is always positive or negative. Evolution then leads to a continuous increase or decrease of *a*_1_, resulting in specialisation (Fig. 3d, A1d,f). When differentiating the fitness function (eq. 3) to get the fitness gradient, it can easily be shown that the loss of one prey systematically selects specialisation on the other prey (Fig. 2a). Finally, evolution toward specialisation can happen after the emergence of polymorphism (branching events). Such branching points (convergent and invasible) eventually lead to the coexistence of two specialists, one on each prey (Figs 2d, 3e, A1e). Such branchings occur only for convex trade-offs where the singularity is invasible (eq. 7). Note that these outcomes are not mutually exclusive, as several singularities may coexist for a given set of parameters, yielding complex evolutionary dynamics. For instance, on Fig. 3e, two repellors coexist with a branching point, so that, depending on its initial diet, a predator can evolve to specialisation on the host (if the initial *a*_1_ is high, i.e., higher than the value of the highest repellor), on the alternative prey (if the initial *a*_1_ is low, i.e., lower than the value of the lowest repellor), or to diversification (i.e., a coalition of two specialists as illustrated on Fig. 2d) (Fig. 3e and A1e).

### 2.5. Parasites affect diet evolution

Now that we have described the possible outcomes, we investigate how the intensity of the parasite-induced virulence and facilitation effects constrain their occurrence. Virulence effects are depicted on Fig. 3, facilitation effects on Fig. A1 and both effects investigated concurrently on Fig. 4. Data were generated with Mathematica®, using analytical form of the fitness gradient and the equilibriums of the four species. For each pair of parameters, we are looking for the roots of gradient fitness. For each, we controlled the existence of all species, and we numerically determined convergency and invasibility.

**Figure 4.**
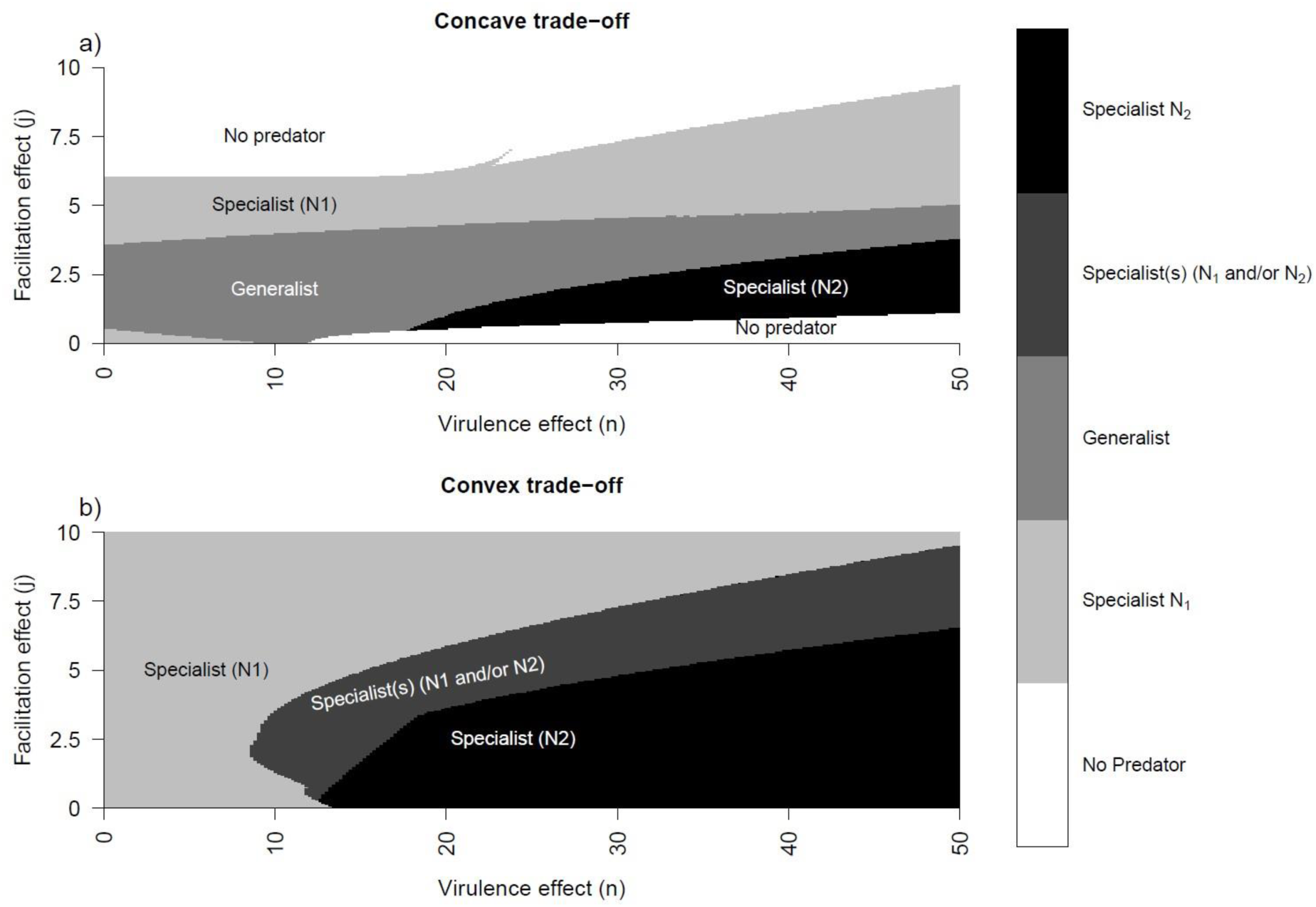
Predator’s diet after evolution as a function of virulence effect (x-axis) and facilitation effect (y-axis) of the parasite, with concave (a) and convex (b) trade-offs. Parameter values: see Table 1, except a) s = 2, b) s = 0.8.

When virulence increases (Fig. 3), as expected, it changes the outcome of evolution from specialisation on the infected prey to specialisation on the alternative prey. Conversely, but also in agreement with our predictions, higher facilitation effects eventually lead to specialisation on the infected prey (Appendix A). In both cases, intermediate effects either lead to generalist strategies (concave trade-offs, Fig. 3b, A1b), or to diversification (convex trade-offs, Fig. 3e, A1e). To go further and assuming concave trade-offs, increasing the virulence effects move the CSS toward lower values of *a*_1_, so that the predator increasingly specialises in the alternative prey (Fig. 3a-c). At the opposite, increasing facilitation moves the CSS toward higher values of *a*_1_ so that the predator increasingly relies on the host species (Fig. A1a-c). Similarly, for convex trade-offs, higher virulence effects move the system from a complete specialisation on the host (Fig. 3d) to a situation in which most dynamics would lead to specialisation on the alternative prey (fig. 3f). Analogous variations can be described when decreasing the facilitation effects (fig. A1d-f).

Figure 4 summarizes these antagonistic consequences of virulence and facilitation effects, and how they depend on the shape of the trade-off function. As previously described, specialist strategies are selected when either virulence or facilitation effects are strong (Fig. 4, light grey and black areas), regardless of the trade-off shape. However, most parasites are likely to induce virulence and facilitation effects simultaneously, so that the evolutionary outcome may often be in between these extreme cases (i.e., in the middle of the panels of Fig. 4). Such evolutionary outcomes depend on the shape of the trade-off. Concave trade-offs allow generalist strategies (medium grey, Fig. 4a) while convex trade-offs favor the diversification of predator diets, potentially leading to the coexistence of specialist strategies (dark grey, Fig. 4b). Finally, note on Figure 4 that extreme effects may lead to the loss of coexistence within the module (white areas), highlighting the fact that evolutionary responses of predators to prey infection may have far reaching consequences for community structure.

### 2.6. Consequences on coexistence

While we know that parasitism may directly constrain prey coexistence by changing the relative weight of direct and apparent competition (Prosnier et al., 2018), here we highlight that evolutionary dynamics in response to parasitism may favor coexistence in trophic modules. To clarify the role of evolution, on figure 5, we show the coexistence conditions with (grey area) and without (area delimited by the solid line) diet evolution. For the “without evolution” scenarios, we consider as reference cases in which the predator is specialised on the infected prey (Fig. 5a), on the alternative prey (Fig. 5c), or is a generalist (Fig. 5b).

**Figure 5.**
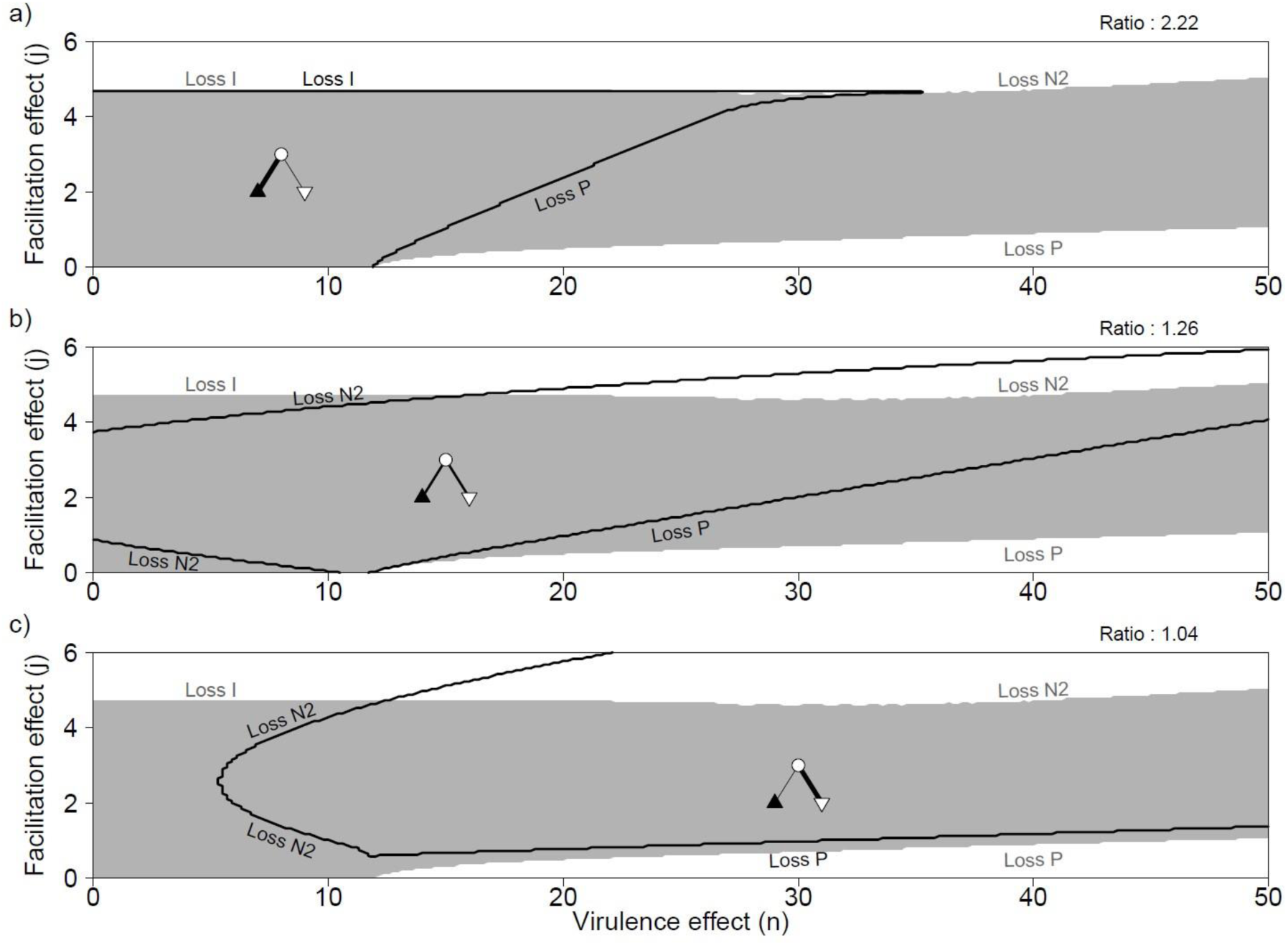
Coexistence of the three species, with convex trade-off, and with (grey area) or without evolution (inside the black delimitation). When the predator is a) specialist on the infected prey *N_1_* (*a_1_* = 1.6), b) generalist (*a_1_* = 0.7), c) specialist on the alternative prey *N_2_* (*a_1_* = 0). Species loss at the limits are written in dark without evolution, and in grey with evolution. Ratio is the ratio of coexistence surface with/without evolution. See Table 1 for parameter values except *s*=2. Note that Fig B1 shows the results with a concave trade-off.

Our results suggest that evolution systematically enhances the size of coexistence area. On the range of parameters we study here, the coexistence area is increased by a 2.22 factor compared to a predator specialist on the host prey (Fig. 5a), by 1.26 compared to a generalist predator (Fig. 5b), and by 1.04 compared to a predator specialist on the alternative prey (Fig. 5c). Evolution particularly acts at low and high values of virulence effects. At low values where the combination of predation and competition could result in the exclusion of the alternative prey (Fig. 5c), evolution allows coexistence because it selects predator diets that are more specialised on the host prey. Thereby, evolution increases apparent competition on the competitively dominant prey. At high values, the predator is excluded because the parasite decreases the amount of prey available, especially in case of specialisation on the host prey (Fig. 5a). Here evolution promotes coexistence in that it favors the diets more oriented toward the alternative prey. Note that while the global effect of evolution is to enhance coexistence, it harms it in particular contexts. For instance, when both virulence and facilitation effects are high, from an ecological point of view a balance exists between apparent and direct competition among prey and coexistence is possible without evolution (e.g. Fig. 5b,c, see also Prosnier et al. (2018)). Evolution however here increases the weight of apparent competition and results in the exclusion of the alternative prey. While Fig. 5 assumes a concave trade-off, we note that the effects of evolution on coexistence are qualitatively similar for convex trade-offs (see Appendix B).

## 3. Discussion

Parasites likely affect food webs in many ways. Because of their virulence and of the resulting decrease in fecundity, they can constrain the availability of prey populations, thereby leading to resource limitation for predators. Alternatively, parasite-induced phenotypic alterations can strengthen trophic interactions, making infected prey more susceptible to predation. While these ecological consequences of parasitism for food webs have received some attention (Banerji et al., 2015; Buck, 2019; Buck & Ripple, 2017; McNaughton, 1992), parasitism also affects evolutionary dynamics of prey and predator species. Here, our model shows that the evolution of predator diet depends on the type of parasitism effect (virulence or facilitation) and on its intensity. Virulence effects usually lead to the selection of increased predation on the alternative (i.e., non-host) prey, while facilitation effects favor predation on the host species. Such results are consistent with our predictions based on host profitability. Virulence effects reduce profitability whereas facilitation effects increase it. Evolution also favors, in general, coexistence among prey and between predators and parasites.

As expected, virulence and facilitation effects have antagonistic consequences on the evolution of predator diet. Virulence effects, here modelled as decreased host fecundity, lead to specialisation on the alternative prey. Because virulence decreases the host prey population, the first part of the fitness function (eq. 2) is decreased and foraging strategies that focus more on the alternative prey are favored. Conversely, facilitation effects (i.e., increase in host vulnerability) lead to specialisation on the infected prey. Again, turning to the definition of fitness (eq. 2), the first term is increased when facilitation effects are present, so that strategies focusing more on the host species are selected. Consequently, the evolution of predator diet can here be understood in the context of optimal foraging (Charnov, 1976a, 1976b; Emlen, 1966; MacArthur & Pianka, 1966). Interestingly, our model shows that, depending of the parasite effect, predator can prefer infected or healthy species, that can be related to a preference for infected individuals (Brandell et al., 2022; Lopez & Duffy, 2021; Mouritsen & Poulin, 2003) or healthy ones (Gutierrez et al., 2022). This results has immediate implication for the healthy herd hypothesis under which the attack of predators eliminates infected individuals (Richards et al., 2022). Our results suggest that such an ecological process could be unstable from an evolutionary point of view, especially when virulence effects are strong. Predators evolve to become generalists when virulence and facilitation effects have intermediate intensities and/or when they act simultaneously. Thus, we expect diet to shift progressively from specialism on one prey to the other prey when the two effects vary. Generalism requires balanced virulence and facilitation effects and concave trade-offs. It is replaced by a coalition of two specialists under convex trade-offs. Such a link between trade-off shape and diversification is consistent with previous theoretical works on the evolution of specialisation (Egas et al., 2004; Rueffler et al., 2006; van Velzen & Etienne, 2013). Note that we have not tested a third scenario in which infection increases the energetical value *e* of the infected prey, as observed in *D. magna* infected by DIV-1 (Prosnier et al., 2024). In this case, we expect a similar result to the facilitation effect, but with a larger coexistence due to smaller predation on infected individuals, thus a smaller risk of losing the parasite.

The evolution of predator diet influences coexistence between the two prey species, because it depends on the balance between direct and apparent competition. It also influences the predator-parasite coexistence, by modulating the resource overlap between the two species. Concerning prey coexistence, virulence effects reduce host competitivity and may ultimately lead to its exclusion (Prosnier et al., 2018). With evolution, however, a decrease in host density selects a lower consumption by the predator, which balances the decrease in host competitive ability and thereby promotes its persistence. Consequently, if we consider the example of DIV- 1, that reduce host competitivity (Decaestecker et al., 2015) due to a high virulence effect (Prosnier et al., 2024), then we may expect that predator diet evolution will ultimately reduce predation on *D. magna*. Similarly, if the alternative prey were to become rare, evolution would select diets that are more focused on the host prey, so that coexistence would again be favored. In a nutshell, adaptive foraging here creates a negative frequency-dependence that promotes the persistence of the rare prey, a mechanism already pointed out by several works (Kondoh, 2003; Křivan & Eisner, 2003; Loeuille, 2010) but not to our knowledge in the context of parasitism. Conversely, when considering facilitation effects, evolution selects higher predation on the host, so that the negative effects of parasitism on prey coexistence are magnified (i.e., both parasitism and evolution increase predation on the host prey).

We observe an increase in predator-parasite coexistence due to niche partitioning between parasite and predator when parasites induce virulence effects. This is consistent with the approaches that unify parasites and predators under the same natural enemy concept (Raffel et al., 2008). Here, the parasite and the predator can be viewed as intraguild predators since they compete for the same trophic resource (i.e., the host population) while the predator also kills the parasite through concomitant predation (i.e., when it consumes infected individuals) (Sieber & Hilker, 2011). Niche partitioning between intraguild predators due to evolutionary dynamics has already been shown (Ingram et al., 2012). While the widespread co-occurrence of parasites and predators in food webs (Lafferty et al., 2006) can be puzzling given the competition that exists between the two groups, our results on the role of diet evolution may offer a possible explanation. Stewart et al. (2018) described what could be interpreted as niche partitioning between the parasites and predators of *Ceriodaphnia* in Lake Gatun (Panama). However, direct comparison with our theoretical work is difficult as the evolutionary processes remain unknown in this empirical system. An empirical validation of our results would involve experimental works based on short-lived organism to allow rapid evolution. For instance, that could be the system used by Banerji et al. (2015): *Paramecium caudatum* infected by *Holospora undulata* and consumed by a rotifer, *Didinium nasutum*, which allows ecological dynamics in less than two months. We could add a competitor to *P. caudatum* through another species of *Paramecium* or another ciliate like *Stylonychia pustulata* as done by Gause (1934). From a theoretical point of view, an interesting development would be a more systematic study of the links between predation and parasitism spread. While predators are often supposed to limit disease spread (see the healthy herd hypothesis mentioned above), positive effects of predators on infection rates are also possible. Indeed, as showed by studies since a few years, some predators, called “predator-spreader”, can positively affect parasites (Richards et al., 2022, 2023). For instance, a predator acts as a spreader when it releases parasites in feces, when it acts as a vector, when it reduces prey immune response due to stress, or when it affects prey behaviour. It is also possible when predators prefer healthy prey, that is similar to the niche separation observed here. We believe that such situations may strongly change the evolutionary dynamics we uncover here, as it creates a direct link between predator and parasite niches.

In this work, we show how parasites may affect the evolution of predator diet and how in return adaptive foraging alters species coexistence. How such relationship between evolution of foraging and parasitism affects more complex food webs would be a natural extension of the present work. For instance, while coexistence of predators and parasites may not be easily explained from an ecological point of view (due to competition), our results suggest that evolution can provide the niche partitioning needed to allow the maintenance of their diversity and could explain why “a healthy system […] is rich in parasite species” (Hudson et al., 2006). However, at this point, it seems premature to extrapolate our results to complex food webs, as empirical information is lacking for several important aspects.

Because we aimed at studying a variety of evolutionary scenarios (flexible trade-offs, different effects of parasitism), we voluntarily simplified ecological dynamics. For instance, we use a linear functional response (close to type-I). While type-II functional responses are sometimes preferred, it is unclear, in the case of parasite-induced facilitation effects, which parameters of the functional response would be modified (i.e., handling times and/or attack rates). Moreover, we expect our main results concerning the effect of adaptive foraging on species coexistence and predator-parasite niche separation to be robust. For instance, considering a type II functional response, it is easy to show that when the infected prey is rare as a consequence of infection, evolution will still favor the consumption of the alternative prey, thereby promoting prey coexistence and niche separation, as with the type I functional response. These deductions are consistent with literature suggesting that adaptive foraging creates a frequency-dependent process that is utterly stabilizing, regardless of the functional response used (Loeuille, 2010; Valdovinos et al., 2010). Finally, studies with a type-I functional response should be considered ecologically relevant according to reality of predation dynamics and prey densities in empirical ecosystems: for instance due to the size ratio, or because prey density is generally low, thus functional response is more in the linear part that in the saturating one (Novak et al., 2025; Sjöberg, 1980).

Importantly, we did not address the cases in which the parasite is trophically transmitted to the predator, which is part of the cycle (see Fleischer et al. (2021) for an example of predator evolution in response to trophically-transmitted parasites). Considering our results, if parasites have also a virulent effect on the predator, then we expect that our results of an evolved niche separation would be magnified, because the parasite would become both a competitor and a parasite for the predator. However, in some well-known cases niche separation is not possible, for instance when the predator is the final host in a complex life cycle. Because such parasites need to infect the predator to complete their cycle, we can deduct from our results some constraints on the virulence and facilitation effects due to the evolution of predator diet. Virulence should be lower both for the prey and the predator (see the model of Kuris (2003), for a low virulence for the predator, but not for the prey, and the review of Fayard (2020) for a low virulence on the intermediate host). However, if avirulence is not possible (due to evolutionary or biological constraints), then the parasite should maximize facilitation whereby the intermediate host is eaten by the final host before its death. Because predation on host is necessary for the parasite, the niche overlap should maintain coexistence between the predator and the parasite, contrary to our system where such situations would lead to parasite extinction. Moreover, it should be beneficial to the predator (Øverli & Johansen, 2019). Consequently, we show from our work and the resulting discussion, that studying facilitation effects is also interesting for non trophically-transmitted parasites, as well as addressing virulence effects for trophically-transmitted parasites.

Finally, despite the simplification inherent in modelling, the possible generalization of our results and the various scenarios investigated allow us to consider possible effects of parasitism in food webs that are currently understudied. Our study may be used as a basis to further empirical works. Promising empirical frameworks exist, as illustrated by the DIV1-*Daphnia magna* system we discussed at several instances, or when considering stickleback species that feed on benthic and/or pelagic prey with varying in infection levels (Fleischer et al., 2021). We believe that combining theoretical developments will empirically testing on the interplay between parasitism and foraging will ultimately allow a more integrative understanding of coexistence constraints within ecological networks.

## Acknowledgments

The authors thank D. Claessen, T. Spataro, and O. Kaltz, for helpful discussions, and D. I. Bolnick and two anonymous reviewers for their comments.

## Data, script, code, and supplementary information availability

Data, script and code are available on Zenodo (Prosnier et al., 2026): https://doi.org/10.5281/zenodo.23459810.

Supplementary information is available after the references:

A. Diet evolution due to parasitism: facilitation effect
B. Consequences on coexistence: concave trade-off

## Supplementary information

### **A)** Diet evolution due to parasitism: facilitation effect

When only facilitation effects occur (Fig. A1), high values always lead evolution to shift from specialisation on the alternative prey to specialisation on the infected prey, which is opposite to virulence effects. Intermediate values either leads to generalist strategies (concave trade-offs, A1b), or to diversification (convex trade-offs, A1e). To go further and assuming concave trade-offs, increasing the facilitation effects move the CSS toward higher values of *a*_1_, so that the predator increasingly specialises on the infected prey (Fig. 1Aa-c). Similarly, for convex trade-offs, higher virulence effects move the system from a complete specialisation on the alternative prey (Fig. A1f), to a situation in which most dynamics would converge to diversification (i.e., the coexistence of two specialists, Fig. A1e), eventually leading to situations in which almost all initial diets will lead to specialisation on the host (fig. A1d). However, note that, in some case, as on Fig. A1d if you start with 0.1<*a*_1_<1 you may loss coexistence of the system through the loss of the parasite (evolution leads to a grey area).

**Figure A1.**
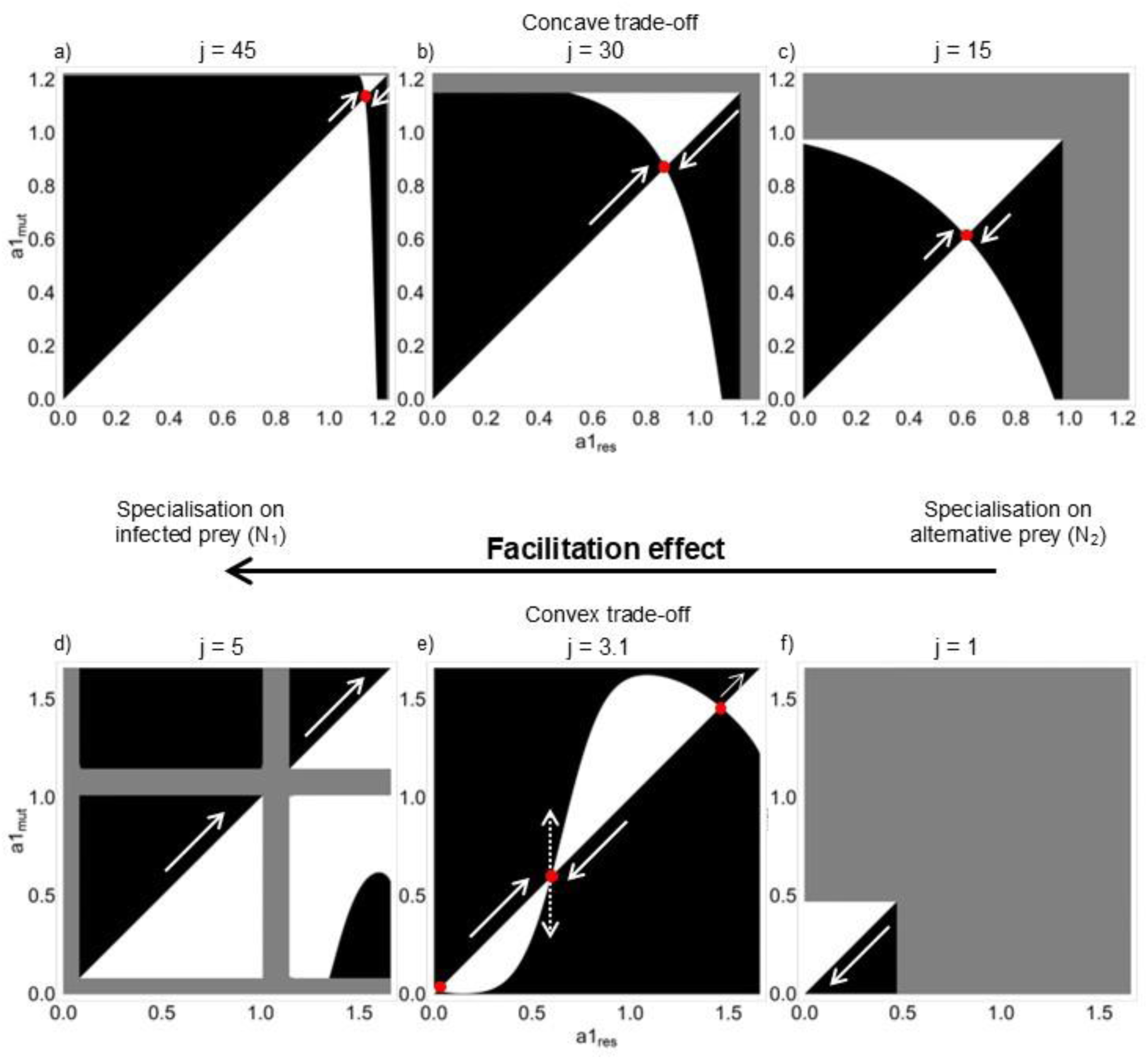
Pairwise Invasibility Plots showing the evolution of predator diet (i.e., evolution of *a_1_*) for concave (a-c) and convex (d-f) trade-offs, when increasing the intensity of the facilitation effect, *n*. Black areas correspond to a positive relative fitness of mutants, white areas to a negative relative fitness of mutants, and grey areas shows no-coexistence of the three species. The white solid arrows show the direction of evolution, the white dotted arrows show evolutionary branching. a-c) CSS, d) no singularity, e) one EBP and two repellors, f) no singularity. Parameter values: see Table 1, except a) *n* = 20, *j* = 45, *s* = 2; b) *n* = 20, *j* = 30, *s* = 2; c) *n* = 20, *j* = 15, *s* = 2; d) *n* = 15, *j* = 5, *s* = 0.8; e) *n* = 15, *j* = 3.1, *s* = 0.8; f) *n* = 15, *j* = 1, *s* = 0.8. Note that Fig 3 shows the results with the virulence effect.

### **B)** Consequences on coexistence: concave trade-off

Similarly, as for convex trade-off, we compare the coexistence conditions with (grey area) and without (area delimited by the solid line) diet evolution. For the “without evolution” scenarios, we consider the cases where the predator is specialised on the infected prey (Fig. B1a), the alternative prey (Fig. B1c), or is a generalist (Fig. B1b).

Our results show that coexistence with evolution is higher than coexistence without evolution. The coexistence area is increased by a 3.58 factor compared to a predator specialist on the host prey (Fig. B1a), by 3.46 compared to a generalist predator (Fig. B1b), and by 1.23 compared to a predator specialist on the alternative prey (Fig. B1c). Evolution particularly acts at low and high intensities of virulence effects. At low intensities where the combination of predation and competition could result in the exclusion of the alternative prey (Fig. B1c), evolution allows coexistence because it selects predator diets that are more specialised on the host prey. Thereby, evolution increases apparent competition on the prey that is, without evolution, competitively dominant. At high intensities, the predator might be excluded because the parasite decreases the amount of prey available, especially in case of specialisation on the host prey (Fig. B1a). Here evolution promotes coexistence in that it favors the diets more oriented toward the alternative prey.

**Figure B1.**
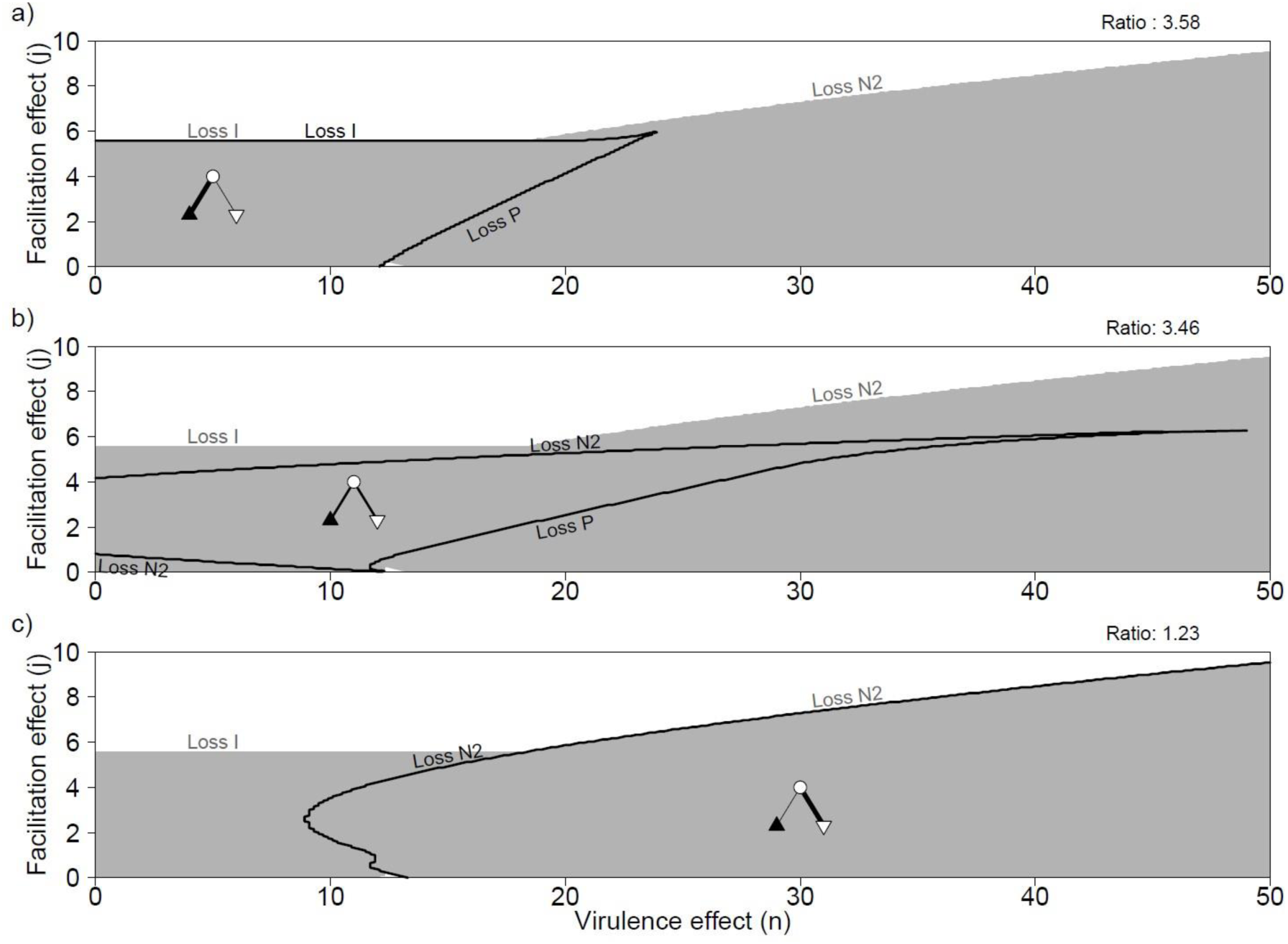
Coexistence of the three species, with concave trade-off, and with (grey area) or without evolution (inside the black delimitation). When the predator is a) specialist on the infected prey *N_1_* (*a_1_* = 1.1), b) generalist (*a_1_* = 1.0), c) specialist on the alternative prey *N_2_* (*a_1_* = 0.2). Species loss at the limits are written in dark without evolution, and in grey with evolution. Ratio is the ratio of coexistence surface with/without evolution. See Table 1 for parameter values except *s*=0.8. Note that Fig. 5 shows the results with a convex trade-off.

